# MELISSA: Semi-Supervised Embedding for Protein Function Prediction Across Multiple Networks

**DOI:** 10.1101/2023.08.09.552672

**Authors:** Kaiyi Wu, Di Zhou, Donna Slonim, Xiaozhe Hu, Lenore Cowen

## Abstract

**Motivation:** Several popular methods exist to predict function from multiple protein-protein association networks. For example, both the Mashup algorithm, introduced by Cho, Peng and Berger, and deepNF, introduced by Gligorijević, Barotand, and Bonneau, analyze the diffusion in each network first, to characterize the topological context of each node. In Mashup, the high-dimensional topological patterns in individual networks are canonically represented using low-dimensional vectors, one per gene or protein, to yield the multi-network embedding. In deepNF, a multimodal autoencoder is trained to extract common network features across networks that yield a low-dimensional embedding. Neither embedding takes into account known functional labels; rather, these are then used by the machine learning methods applied after embedding.

**Results:** We introduce MELISSA (MultiNetwork Embedding with Label Integrated Semi-Supervised Augmentation) which incorporates functional labels in the embedding stage. The function labels induce sets of “must link” and “cannot link” constraints which guide a further semi-supervised dimension reduction to yield an embedding that captures both the network topology and the information contained in the annotations. We find that the MELISSA embedding improves on both the Mashup and deepNF embeddings in creating more functionally enriched neighborhoods for predicting GO labels for multiplex association networks in both yeast and humans.

**Availability:** MELISSA is available at https://github.com/XiaozheHu/melissa

## Introduction

In 2016, Cho et al. introduced their groundbreaking Mashup algorithm for function prediction by integrating information across multiplex biological networks [7]. Mashup consists of three steps: in the first step, the Random Walk with Restart (RWR) algorithm [23] is run to produce a diffusion state vector separately for each node for each network. In the second step, a low-dimensional embedding is constructed to minimize the distance to all the individual network-specific vectors globally. In the last step, these global low-dimensional feature vectors are then passed to classifiers such as *k*-nearest neighbors [6] or support vector machines [3] in order to accomplish the functional label prediction.

However, in the Mashup paradigm, we notice that biological knowledge, encoded in the form of the GO [8] functional labels on the training set nodes, is only incorporated in the last step of this process. The embedding itself, constructed in the first and second steps, is entirely unsupervised and comes only from the topological structure of the networks. The same is true of deepNF [14], an alternative method that constructs the embedding using a multi-modal deep autoencoder. Therefore, the motivation of this paper is to incorporate information from the known GO labels of the nodes in the training set into the low-dimensional embedding to improve its quality and, thus, improve on the state-of-the-art Mashup and deepNF algorithms.

One popular approach to including such biological label information is *semi-supervised* graph embedding methods [28, 1, 2, 26]. The biological information, in the form of functional labels, can give rise to a set of “must-link” (ML) constraints and a set of “cannot-link” (CL) constraints [1]. These constraints then guide the embedding procedure toward a result that encodes both the inherent structure of the data as well as the information contained in the functional annotations.

In the protein function domain, genes have multiple functional labels, both noisy and incomplete [10]. So it can be challenging to generate both types of constraints. The incompleteness is a challenging problem: “cannot link” constraints require proteins known *not* to be involved in some function. But in our setting, the fact that a gene lacks a particular functional annotation does not necessarily indicate that the gene does not play a role in that function. It may very well be that it simply has not been experimentally observed yet.

One possible way to handle the above challenges, on a gene-by-gene basis, would be to use some of the sets of specially curated negative GO annotations that the community has begun to construct [12, 25]. Our method, MELISSA, instead takes a different approach. We augment each network with a sparse set of artificial new nodes, which also are involved in the embedding step. Intuitively, the new artificial nodes represent the “centers” of a coarse clustering of functional labels. We place ML constraints between the original nodes in the training set and each new artificial node that contains one of its GO functional labels to bind genes in the training set to their cluster label and encourage them to cluster. At the same time, CL constraints are placed between the artificial nodes themselves to encourage the clusters to separate from each other (See Section 2.4). MELISSA uses a biclustering procedure [9] on the annotation matrix that maps genes to GO labels of appropriate specificity to generate the set of coarse cluster labels that will be assigned as artificial cluster nodes.

After augmentation, the resulting networks now contain positive (ML) and negative (CL) weights. The standard RWR approach to compute each node’s diffusion state vectors cannot be directly applied. Thus, in MELISSA, we adopted a signed version of the graph Laplacian [16, 18, 13] and generalize the diffusion state representations of each node to the networks with both positive and negative weights. Then, the rest of the Mashup pipeline, from dimension reduction to function prediction based on the low-dimensional embedding, proceeds as before.

Once the embedding is formed, a variety of different classification methods can be applied to the embedded space. Because our focus is on improving the information content of the embedding, in this work we pair MELISSA with the simple *k*NN classifier, where comparing functional label prediction of competing embeddings gives a sense of the functional enrichment in local neighborhoods. As shown below, in this setting, MELISSA improves the overall performance of the functional label prediction task, compared to the original Mashup and the deepNF embeddings, demonstrating its ability.

Thus MELISSA can be used to analyze multiple networks organized by a guilt-by-association property and provide an accurate and scalable framework for network integration and analysis from different experiments.

## Methods

MELISSA varies from Mashup by including a step of network augmentation to encode functional information before the embedding and the learning phase. This step requires augmenting the networks with auxiliary cluster nodes which we induce from the available gene annotations. A summary of the procedure is given in Figure 1.

**Fig. 1:**
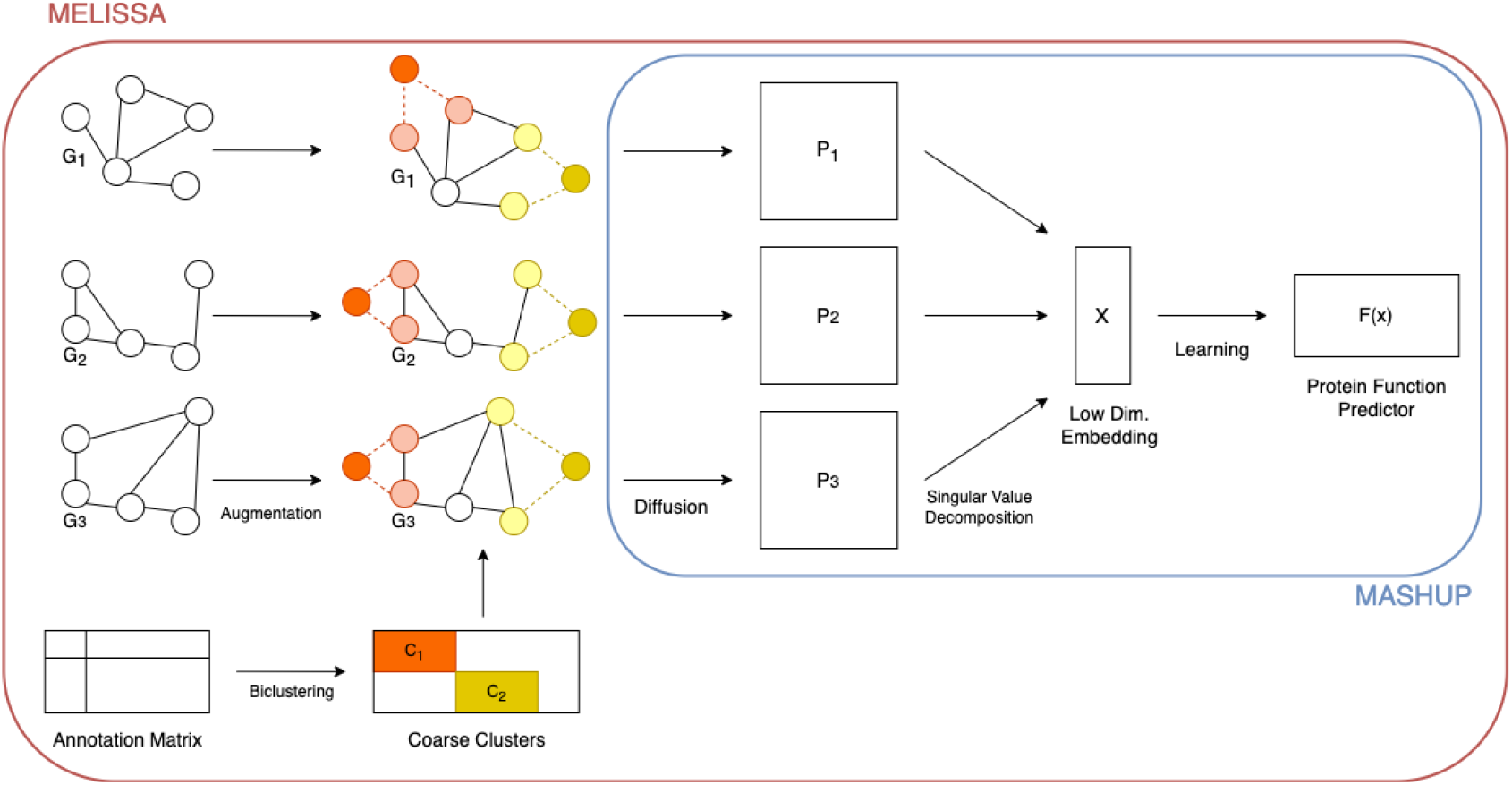
Workflow of MELISSA extending Mashup. Nodes and labels are first biclustered, and each network is augmented with one auxiliary node for each functional label cluster. ML constraints (dashed edges) are added to pull the nodes in the same cluster closer to each other, while CL constraints (solid edges) are added to push the clusters away from each other in the embedding. Then we run the Mashup procedure as summarized in Section 2.2 on each of the augmented networks.

### Preliminaries and Notation

The datasets that we consider consist of a collection of networks *G*^*i*^ = (*V, E*^*i*^, *w*^*i*^) with *i* ∈ *{*1, …*N}* which share a set of nodes *V* but each has its own set of edges *E*^*i*^ and the set of edge weights *w*^*i*^ *>* 0. In our work, the nodes correspond to genes that appear in the union of all the networks. The edges *E*^*i*^ correspond to a different type of relationship between pairs of genes for each network. As in the experiment in the original Mashup paper, these relations range from experimental evidence of interaction or association between two genes, to co-expression, among other things (see [7]). Finally, the weights *w*^*i*^ indicate the confidence in the edges being correct.

The adjacency matrices of the graph *G*^*i*^ are denoted by ***A***^*i*^, and their probability transition matrices are denoted by ***T*** ^*i*^ where 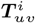 denotes the probability that a random walk on graph *G*^*i*^ at vertex *v* transitions to *u* in one step. The functional annotations of the genes are represented by the binary matrix ***B*** ∈ *{*0, 1*}*^*l×n*^ where *l* is the number of distinct labels and *n* = |*V* | is the number of the genes. In ***B***, each column corresponds to the set of annotations a gene has been given, and each row corresponds to the set of genes with a given annotation.

### Review of the Mashup Embedding

The original Mashup procedure consists of the following three core steps:

1. **Diffusion**. On each *G*^*i*^, a diffusion process is run which creates a RWR matrix representation ***W*** ^*i*^ ∈ ℝ^*n×n*^ of the network.
2. **Embedding**. A shared embedding is created using the matrix representations generated in the diffusion step. This is achieved via a singular value decomposition or dictionary learning techniques. Ultimately this gives a *d*-dimensional vector representation of every node in the dataset.
3. **Learning**. Once every node in the dataset is represented by a vector, existing function prediction methods can be applied using the embedding and the available annotations. The original Mashup uses a support vector machine (SVM) for the final learning step. However, it also carefully filtered the set of GO labels that it considered to discard GO labels whose set of labeled genes overlapped (with Jaccard threshold greater than .1), advantaging methods such as SVM that search for a separating hyperplane. Since we are also interested in the less artificial setting where GO labels are allowed to overlap, we instead use *k*NN as the classifer for results in this paper.

Regardless of which classifier is used at the end of the pipeline, we consider the **Diffusion** and **Embedding** steps to be the main contribution of Mashup and briefly describe them in detail.

The **Diffusion** step generates a matrix representation of each network. There are many ways of generating node embeddings, such as Diffusion State Distance [6, 5], Node2vec [15], and spectral methods [19]. Mashup adopted a Random Walk with Restart (RWR) based approach [17], i.e., for each vertex *u* ∈ *G*^*i*^, Mashup iteratively computes the *t*-step RWR distribution 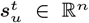 as follows, let 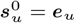,

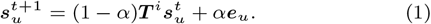

where ***e***_*u*_ ∈ ℝ^*n*^ is the vector with entry 1 at the *u*-th index and 0 elsewhere and 0 ≤ *α <* 1. Following [6, 5, 7], the node embedding of *u* is the *diffusion state* 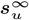, the stationary distribution at the fixed point of the iteration (1). By stacking the diffusion states 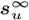 as columns, we obtain the diffusion state matrix ***W*** ^*i*^, which is the RWR representation matrix used in Mashup.

Once each set of *N* networks has been represented by its associated RWR matrix ***W*** ^*i*^, Mashup then combines the matrix representations and constructs a low-dimensional embedding. The original Mashup framework proposed two embedding strategies, the first being a dictionary learning approach and the second being the singular value decomposition (SVD). For the sake of simplicity, we focus on the SVD approach, i.e., the low-dimensional embedding ***X*** ∈ ℝ^*d×n*^ are formed by the scaled largest *d* left singular vectors of the concatenated matrix log ***S*** where ***S*** = [(***W*** ^1^)^*T*^, *…*, (***W*** ^*N*^)^*T*^]^*T*^ and log(*·*) denotes the element-wise logarithm. As suggested in [7], for the optimization purpose in the implementation, the SVD-based embedding can be computed by the eigenvalue decomposition of the *n × n* matrix 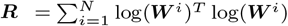. A small constant (e.g., reciprocal of the number of genes) was added to each entry of ***W*** ^*i*^ to avoid taking the log of zero entries. Taking the top *d* eigenvalues **Λ** = diag(*λ*_1_, *…, λ*_*d*_) and eigenvectors ***U*** = (***u***_1_, *…*, ***u***_*d*_) ∈ ℝ^*n×d*^, the low-dimensional embedding is ***X*** = **Λ**^1*/*4^***U*** ^*T*^.

Note that Mashup as described in [7] involves concatenating the diffusion state matrices from different networks vertically when performing the joint factorization; the Mashup code allows us to concatenate either horizontally or vertically. In our experiments, we found the vertical approach always performed better than the horizontal one.

Our key observation is that Mashup uses only network topology to construct its embedding. Our objective is to attain a more functionally meaningful low-dimensional embedding of the networks by augmenting the original networks so that the **Diffusion** and **Embedding** steps are aware of known functional annotations. We hypothesize that the improvements in the embedding transfer to the performance in the **Learning** step.

### Review of the deepNF Embedding

DeepNF [14], introduced in 2018, tries to learn a useful low-dimensional embedding of proteins with a multimodal deep autoencoder (MDA) [24] that preserves non-linear network structure across multiple networks characterized by diverse connectivity patterns. In [14] it is shown that deepNF preserves the non-linear network structure with its deep neural network (DNN) architecture in an efficient and scalable manner, and at the same time denoises the links in the networks.

The deepNF method involves the following three steps:

1. **Pre-processing**. On each network *G*^*i*^, a RWR procedure is run to create its matrix representation ***W*** ^*i*^ ∈ ℝ^*n×n*^. Then each RWR matrix is converted into a Positive Pointwise Mutual Information (PPMI) matrix ***Q***^*i*^ ∈ ℝ^*n×n*^ that captures the structural information of the network.
2. **MDA Embedding**. A MDA is trained that takes the PPMI matrices as input. A canonical *d*-dimensional feature representation across the networks is extracted from the middle layer of the MDA.
3. **Learning**. The middle layer of the MDA which serves as the low-dimensional vector representation of every node in the networks is then fed into function prediction classifiers.

We elaborate on each step in a bit more detail next. In the **pre-precessing** step, each RWR matrix ***W*** ^*i*^ is generated in the same way as in the Mashup **Diffusion** step. After that, the PPMI matrix ***Q***^*i*^ for the *i*-th network is computed as,

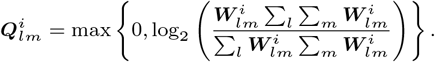

Once each network has been transformed into its information-rich matrix representation, deepNF integrates the PPMI matrices with MDA to construct a low-dimensional feature representation that best approximates all networks. In particular, deepNF first trains a low-dimensional non-linear embedding for each biological network and then concatenates the network embeddings from the previous step into a single hidden layer, allowing the MDA to learn feature representations using all networks. The single bottleneck layer of the MDA is then extracted as the integrated low-dimensional feature representation. Mini-batch stochastic gradient descent with momentum is used to train the MDA.

### Semi-Supervised Embedding via Graph Augmentation

Although the networks *G*^1^, *…, G*^*N*^ have edges that encode interactions and can be used to reconstruct protein functions, we intend to fully utilize any known protein functions from the beginning, even before the **Diffusion** step starts. This can be done by augmenting the original networks using the ML and CL constraints and employing a semi-supervised embedding approach. However, directly adding the ML and CL constraints between the original nodes might outweigh the original edges and, as a result, destroy the original networks’ clustering structure. Therefore, we first augment the networks with auxiliary nodes that encode functional information and then apply the constraints to gently enhance the clustering structure without polluting it.

To introduce the auxiliary nodes that encode functional information, we simultaneously bicluster the proteins and the function labels (see Figure 2). Within the resulting biclusters, each pair of proteins has similar function labels, and functional labels are rarely shared across clusters. Therefore, this suggests we introduce one auxiliary node for each label cluster. There are several popular algorithms for producing biclusters [20]. MELISSA uses the method of [9], because it tends to produce more balanced biclusters, resulting in better practical performance. The number of biclusters is a parameter of MELISSA. We tested setting the number of biclusters to 2, 4, 6, 10, 20 and 40 on 9 different sets of GO labels in the yeast data allowing overlapping GO labels, separating out GO terms of three different frequencies over the three different GO hierarchies (BP, MF and CC). We found that while the best setting varied, it was never useful to have more than 10 biclusters. Results of this exploration appear in Figure 4. Furthermore, 2 biclusters performed reasonably well across all experiments, so in what follows, we report MELISSA results for both yeast and human networks using 2 biclusters.

**Fig. 2:**
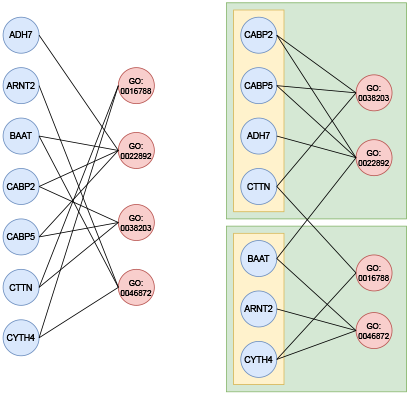
Left: A fragment of the bipartite graph with gene nodes connected to their labeled GO terms, with nodes ordered lexicographically. Nodes on the right are: GO:002678 hydrolase activity; GO:0022892: transmembrane transporter activity; GO:0038203 TORC2 signalling; and GO:0046872 metal ion binding. Right: The same graph with nodes and labels grouped by biclustering, where the transmembrane transporter activity and metal ion binding labels are placed in the same bicluster, with a separate bicluster containing the TORC2 signaling label and the hydrolase activity. The biclusters are highlighted in green. The nodes BAAT and CTTN will be associated with both biclusters because of the cross edges.

**Fig. 3:**
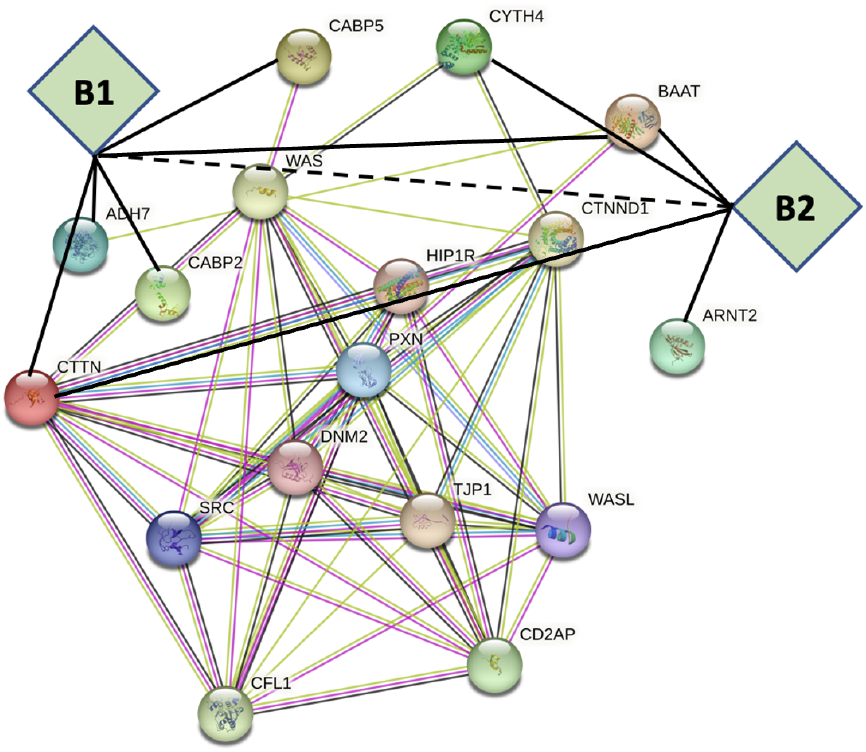
A subset of the augmented STRING network with the two new artificial bicluster nodes from the Figure 2 example added. In particular, B1 represents the top bicluster and B2 represents the bottom bicluster in the Figure 2 example. Solid lines represent the added must-link constraints with positive weight, and the dashed line represents the “cannot link” constraint that attempts to pull the biclusters apart. These constraints help tighten the clustering for the labeled data, but simultaneously pull the labeled data away from unlabeled data in the embedding, making the strength of the edges to the artificial nodes crucial to determining the success of the approach.

**Fig. 4:**
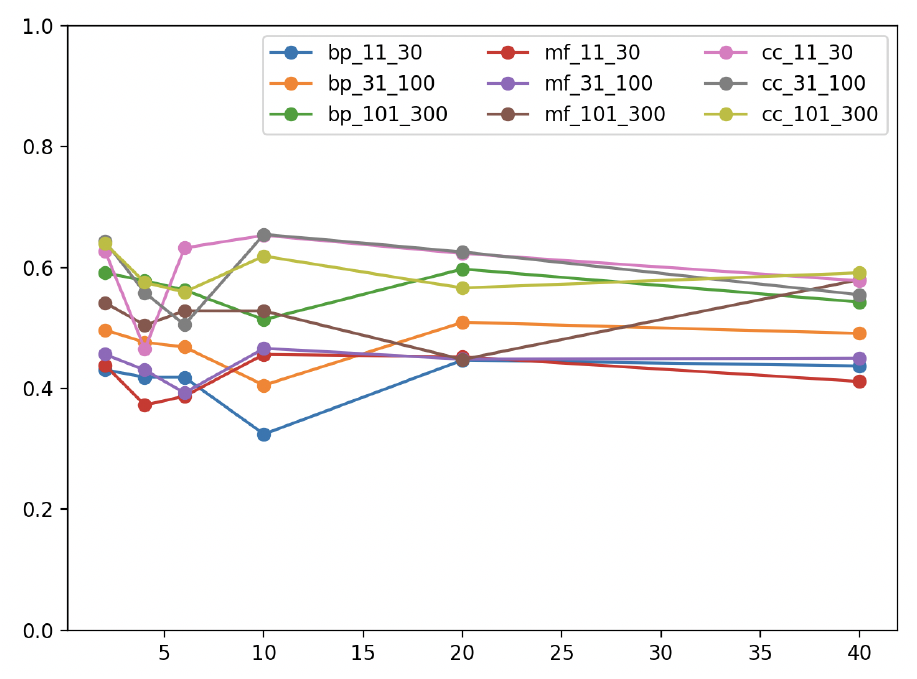
Function prediction comparison with different bicluster numbers for overlap included yeast data. The functional labels used are GO labels in the BP, MF, and CC hierarchies. The x-axis is the bicluster number, and the y-axis is the prediction accuracy.

The next step is to augment the graphs by introducing one auxiliary node for each label cluster and adding the ML and CL constraints as suggested in [27]. More precisely, in each network, we link each protein to the auxiliary nodes that represent the core of the function label clusters according to the protein’s functional labels and put positive weight on the added edges to pull the nodes in the same cluster close to each other. Those added edges are the ML constraints and we denote its set by *E*_ML_ with weight *w*^+^ *>* 0. In addition, we add pairwise CL constraints between the auxiliary nodes to push the clusters away from each other in the embedding. The set of those added edges is denoted by *E*_CL_ with weight *w*^−^ *<* 0. Note, for the sake of simplicity, we drop the superscript *i* in this section.

Since the augmented networks become *signed* graphs due to the negatively weighted edges introduced by the CL constraints, we need to look at the so-called *signed* graph Laplacians [16, 18, 13] to understand the effects of the added ML and CL edges. Let 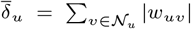 be the *signed* weighted degree of node *u*, where *𝒩*_*u*_ denotes the neighboring nodes of node *u*, and 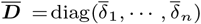 be the *signed degree matrix*. Then the *signed graph Laplacian* is defined by 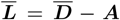. Let ***Y*** ∈ ℝ^*d×n*^ be the *d*-dimensional embedding of the nodes, as usual, the energy function *ℰ* (***Y***) is defined as

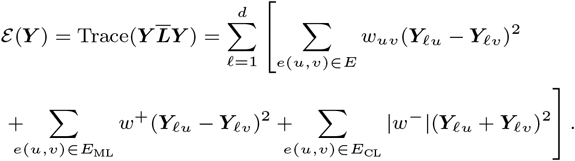

To minimize the energy *ℰ* (***Y***), for *e*(*u, v*) ∈ *E*_ML_, we should have ***Y***_*ℓu*_ ≈ ***Y***_*ℓv*_ which pulls the nodes in one cluster closer to the corresponding auxiliary node and, therefore, the nodes within the same cluster will be placed closer to each other indirectly. On the other hand, for *e*(*u, v*) ∈ *E*_CL_, we expect ***Y***_*ℓu*_ ≈ −***Y***_*ℓv*_, which pushes the auxiliary nodes away from each other and, therefore, places the nodes belong to different clusters apart indirectly.

As we can see, the edge weights *w*^+^ and *w*^−^ for the ML and CL constraints are parameters of MELISSA. In our initial study, since the edge weights on the original networks are confidence scores ranging between (0, 1], we simply set *w*^+^ = 1 and *w*^−^ = −1 in our numerical tests to demonstrate the idea of MELISSA. We plan to optimize those weights in our future work.

Once the augmented graphs are constructed, we still want to use the RWR matrix to represent the graph. In MELISSA, we still use the diffusion state matrix. Thus, we need to generalize the diffusion state matrix ***W*** to signed graphs. For graphs with only positively weighted edges, from (1), by direct calculation, we have, for *α* ∈ (0, 1),

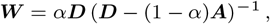

where ***D*** = diag(*δ*_1_, *…, δ*_*n*_) is the weighted degree matrix with 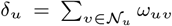 being the usual weighted degree for node *u*. Thus, a natural generalization to a signed graph is obtained by replacing the weighted degree matrix ***D*** with the signed weighted degree matrix 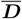 and defining the diffusion state matrix for signed graphs as follows,

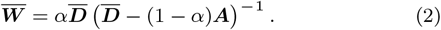

We use this approach to construct 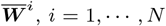, for each augmented graph and then follow Mashup’s steps to combine those matrix representations to obtain a low-dimensional embedding. Due to the presence of the negatively weighted CL links, the entries of 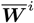 might be negative and, therefore, we increase the small constant used in Mashup to avoid taking the log of negative entries 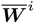. In our experiments, we set this smoothing constant to be the reciprocal of the number of genes plus 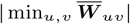. Finally, the low-dimensional embedding of MELISSA is constructed the same as the **Embedding** step of Mashup.

### MELISSA

Our proposed MELISSA procedure consists of five core steps as follows, where steps 3-5 duplicate Mashup with augmented networks.

1. **Biclustering**. Bicluster the annotation matrix ***B*** to obtain coarse groupings of the nodes. The number of biclusters is the parameter *Nc*.
2. **Graph Augmentation** Add new auxiliary nodes corresponding to the function label biclusters. We create two types of new edges involving these auxiliary nodes. First, we add “must-link” edges of weight *w*^+^ that connect each gene to each bicluster that contains one of its matching functional annotations. Then, we add “cannot link” edges of weight *w*^−^ between all pairs of the auxiliary nodes.
3. **Diffusion**. On each augmented graph 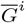 a diffusion process (2) is run which creates a matrix representation 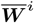of the augmented network.
4. **Embedding** A shared low-dimensional embedding is created using the matrix representations 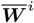. This is achieved via a singular value decomposition or dictionary learning techniques. Ultimately this results in a *d*-dimensional vector representation of every node in the dataset.
5. **Learning**. Once every node in the dataset is represented by a vector, existing function prediction methods can be applied. For example, an SVM approach, like Mashup or DeepNF uses, or a (weighted) majority voting classifier by *k*-nearest neighbors (*k*NN) (similar to [6]) can be trained using the embedding and the available annotations.

## Experimental Setup

### Datasets

Our first experimental setup matches the setup in the original Mashup paper as much as possible. The set of networks we consider in this paper are the ones used in the original Mashup paper. These are from the STRING database v9.1 [11], excluding links derived from text-mining. In particular, we consider six heterogeneous networks over 6, 400 genes with the number of edges varying from 1, 361 to 314, 013 for yeast, and 18, 362 genes with the number of edges varying from 3, 717 to 1, 544, 348 for human. MELISSA, like Mashup, incorporates STRING confidence weights on the edges when computing the diffusion matrix.

The original Mashup paper used GO annotations for human networks and the now-deprecated MIPS annotations for yeast networks. To compare to the original Mashup, we used the same human annotations. For yeast, however, we decided not to use MIPS and instead also consider the GO, so our results are not directly comparable to the results reported in the original Mashup paper for yeast. (Our yeast GO annotations are from the Gene Ontology Consortium [22] (downloaded from FuncAssociate3.0 [4] on 02/12/19)). The GO functional labels are grouped into three distinct functional hierarchies: Biological Process(BP), Molecular Function(MF), and Cellular Component(CC), where we again mimic Mashup to filter the GO terms to only retain those of intermediate specificity, labeling more than 10 and fewer than 301 genes among the nodes. This label set can be further divided into levels of varying specificity, each containing labels that annotate 11-30, 31-100, and 101-300 genes, respectively [21]. We consider 9 different functional annotation experiments, parameterized by one of the 3 hierarchies (BP, MF, and CC) times one of the three levels of specificity of GO terms (11-30, 31-100, and 101-300), and construct MELISSA embeddings for each, see below.

Mashup filters out too similar GO labels within the same level based on Jaccard similarity. We note, however, that the filtering to remove similar GO labels, as in the original Mashup paper, is quite drastic, and we want MELISSA to also work in the typical use case, where GO labels considered do overlap. Below we report results both with (to be most comparable to the original MASHUP) and without GO filtering.

### Evaluation

We compare the performance of the original Mashup and deepNF embeddings with MELISSA in predicting GO functional labels in each of the three hierarchies (BP, MF, and CC) on both the human and yeast multi-network collections in 5-fold cross-validation. Because our focus is on the embedding part, where Mashup and MELISSA vary, we decided to use the computationally less expensive learning method, *k*NN, for function prediction as in [6]. We note that it is possible to achieve better performance than *k*NN with a more sophisticated SVM method [7, 14] (see Discussion section, below).

After obtaining the low dimensional data representation for each node in the test set, we find its *k* nearest neighbors and then cast a majority vote weighted by the reciprocal of the pairwise distance. The label with the most votes is assigned to the gene as its predicted function, and this is considered correct if it matches at least one of its known annotations. The percentage accuracy is the percentage of nodes given correct annotations. We measure this, F1 score, and area under the precision-recall curve (AUPRC) broken down by both hierarchy (Molecular Function (MF), Biological Process (BP) or Cellular component (CC)) and specificity of GO terms (GO terms that label 11-30, 31-100 and 101-300 nodes in the dataset), for both the yeast and human datasets, matching the experimental setup in [7].

In the human network, we chose the same set of GO terms as was chosen by MASHUP, and evaluated on the same sets of GO term range specificities (note that some of the GO terms are obsolete in the current version of the GO). In the yeast network, we switched to the GO (the original MASHUP paper used MIPS), and used an up to date set of GO annotations. For both the yeast and human networks, we evaluate our methods in two settings: 1) using all GO terms, not filtered for overlap, and 2) using the MASHUP procedure to eliminate overlapping GO terms. Note that scenario 2) is evaluating MELISSA on exactly the same set of human GO terms as was considered in the original MASHUP paper.

## Results

For both human and yeast trials, we included Mashup trials with the same setup as presented in the experiments in [7] (including concatenating the diffusion state matrices vertically in the **Embedding** step). We performed five independent runs using five different random seed and all the reported results are given by averaged performance.

We need to set several parameters for the MELISSA framework: the number of biclusters, the embedding dimension, the strength of the ML and CL constraints, and the number of nearest neighbors for majority voting. We set these parameters by comparing MELISSA to the original Mashup (on the original set of GO labels also tested by Mashup). We fixed the number of neighbors for the majority voting at 10, as recommended by [6].

We adopted an embedding dimension of 400 for both yeast and human experiments in both Mashup and MELISSA; this is in line with the guidance of [7] which suggests retaining 5%–10% of the original number of dimensions.

To decide on the number of biclusters, we evaluated different numbers of biclusters on just the yeast data. Figure 4 shows the accuracy using 2, 4, 6, 10, 20, and 40 clusters on the yeast dataset. As cluster number increases, MELISSA’s performance tends to drop slightly, then increase slightly, and then drop again. These results suggest that using just two biclusters does quite well, so we chose two biclusters for both the yeast and human experiments (without any further parameter evaluation on the human data).

For deepNF embedding training, we adopted the suggested parameters from [14]. The MDA architectures are [6 *× n*, 6 *×* 2000, 600, 6 *×* 2000,6 *× n*], and [6 *× n*, 6 *×* 2500, 9000, 1200,9000,6 *×* 2500,6 *× n*] for the yeast and human networks respectively. The stochastic gradient descent optimizer uses batch size of 128 for yeast networks and 256 for human networks, with momentum of 0.95 and learning rate of 0.2 for both yeast and human. The middle layers of the MDA are extracted as embeddings and are then fed into the *k*NN function prediction classifier described above. We ran deepNF with its recommended parameters.

In our experimental setup, where we chose to mimic the experiments run in the original Mashup paper as closely as possible, using two biclusters and allowing both ML and CL constraints seems to do quite well across the different hierarchies and GO-term ranges in both the yeast and human species. Averaged results for five independent runs with these parameters appear in Table 1 and Table 2. Interestingly, this version of MELISSA consistently outperforms both Mashup and deepNF when measured in accuracy, F1 score, or AUPRC score, with the only exception of Yeast CC 101 − 300 trials.

**Table 1.** Function prediction performance on yeast PPI network with overlap removed GO functional labels from the BP, MF, and CC hierarchies based on MELISSA, Mashup, and deepNF data features by majority voting with 10 nearest neighbors in 5-folds cross-validation. The embedding dimension is 400 for Mashup and MELISSA and the MELISSA parameters are: *w*^+^ = 1, *w*^−^ = −1, and 2 biclusters. The deepNF embedding dimension is 600. Results are averaged from 5 independent runs.

| GO<br>HR. | Term<br>Range | Accuracy |  |  | F1 |  |  |
| --- | --- | --- | --- | --- | --- | --- | --- |
|  |  | Mashup | deepNF | MELISSA | Mashup | deepNF | MELISSA |
| BP | 11-30 | 37.91 ± 0.45 % | 36.98 ± 0.64 % | **43.32 ± 0.65 %** | 28.30 ± 0.25 % | 28.71 ± 0.43 % | **31.35 ± 0.48 %** |
| BP | 31-100 | 46.85 ± 0.35 % | 46.45 ± 0.65 % | **50.76 ± 0.48 %** | 34.47 ± 0.25 % | 34.05 ± 0.10 % | **36.11 ± 0.21 %** |
| BP | 101-300 | 59.99 ± 0.46 % | 58.85 ± 0.37 % | **63.45 ± 0.56 %** | 44.55 ± 0.34 % | 43.73 ± 0.15 % | **45.72 ± 0.27 %** |
| MF | 11-30 | 41.47 ± 0.83 % | 42.10 ± 0.54 % | **47.77 ± 0.52 %** | 31.41 ± 0.47 % | 30.46 ± 0.23 % | **34.05 ± 0.30 %** |
| MF | 31-100 | 45.00 ± 0.26 % | 42.57 ± 0.55 % | **46.21 ± 1.01 %** | 34.46 ± 0.24 % | 32.41 ± 0.18 % | **34.85 ± 0.19 %** |
| MF | 101-300 | 56.26 ± 0.64 % | 51.01 ± 0.41 % | **56.90 ± 0.87 %** | 41.98 ± 0.30 % | 40.09 ± 0.37 % | **42.19 ± 0.48 %** |
| CC | 11-30 | 62.74 ± 0.37 % | 63.95 ± 0.42 % | **64.51 ± 1.04 %** | 39.31 ± 0.12 % | **40.74 ± 0.17 %** | 40.08 ± 0.54 % |
| CC | 31-100 | 60.86 ± 0.80 % | 65.54 ± 0.13 % | **66.55 ± 0.55 %** | 41.27 ± 0.14 % | 42.79 ± 0.18 % | **43.87 ± 0.07 %** |
| CC | 101-300 | 71.49 ± 0.27 % | **76.00 ± 0.39 %** | 71.59 ± 0.93 % | 47.07 ± 0.14 % | **47.93 ± 0.20 %** | 45.89 ± 0.80 % |

**Table 2.** Function prediction performance on human PPI network with overlap removed GO functional labels from the BP, MF, and CC hierarchies based on MELISSA, Mashup, and deepNF data features by majority voting with 10 nearest neighbors in 5-fold cross-validation. The embedding dimension is 400 and the MELISSA parameters are: *w*^+^ = 1, CL *w*^−^ = 1, and 2 biclusters. The deepNF embedding dimension is 1200. Standard deviation was calculated from 5 independent runs.

| GO<br>HR. | Term<br>Range | Accuracy |  |  | F1 |  |  |
| --- | --- | --- | --- | --- | --- | --- | --- |
|  |  | Mashup | deepNF | MELISSA | Mashup | deepNF | MELISSA |
| BP | 11-30 | 25.16 ± 0.42 % | 21.44 ± 0.46 % | **28.63 ± 0.13 %** | 18.85 ± 0.11 % | 16.68 ± 0.30 % | **20.61 ± 0.20 %** |
| BP | 31-100 | 30.27 ± 0.35 % | 25.69 ± 0.24 % | **33.62 ± 0.26 %** | 21.85 ± 0.38 % | 19.29 ± 0.11 % | **23.67 ± 0.26 %** |
| BP | 101-300 | 40.64 ± 0.47 % | 34.51 ± 0.23 % | **43.74 ± 0.48 %** | 30.71 ± 0.25 % | 26.21 ± 0.10 % | **32.45 ± 0.23 %** |
| MF | 11-30 | 31.55 ± 0.72 % | 30.28 ± 0.75 % | **37.04 ± 0.29 %** | 24.71 ± 0.36 % | 22.71 ± 0.31 % | **27.63 ± 0.27 %** |
| MF | 31-100 | 33.85 ± 0.33 % | 30.23 ± 0.35 % | **37.11 ± 0.38 %** | 26.74 ± 0.19 % | 23.74 ± 0.35 % | **28.66 ± 0.42 %** |
| MF | 101-300 | 47.47 ± 0.21 % | 33.21 ± 0.25 % | **47.72 ± 0.39 %** | 36.12 ± 0.12 % | 29.69 ± 0.40 % | **37.33 ± 0.14 %** |
| CC | 11-30 | 42.43 ± 1.03 % | 43.56 ± 0.40 % | **48.19 ± 0.48 %** | 29.02 ± 0.31 % | 28.93 ± 0.41 % | **31.44 ± 0.24 %** |
| CC | 31-100 | 46.91 ± 0.22 % | 44.93 ± 0.20 % | **51.11 ± 0.16 %** | 32.55 ± 0.18 % | 30.29 ± 0.35 % | **34.32 ± 0.33 %** |
| CC | 101-300 | 49.51 ± 0.72 % | 45.19 ± 0.45 % | **52.39 ± 0.52 %** | 34.85 ± 0.14 % | 32.58 ± 0.11 % | **36.27 ± 0.14 %** |

However, we note that this experimental setting is suboptimal for assessing biclustering parameters, because Mashup filtered GO terms at each level, removing terms that had Jaccard similarity greater than 0.1 with another category in the same level in order to avoid statistical artifacts arising from overlapping functional categories. Thus we are already looking at a sparser subset of GO terms that are too distinct to form good biclusterings.

We thus repeated the experiments after introducing the overlapping GO terms back into the experimental setting, while keeping all other parameters the same. Averaged accuracy and F1 score across five independent runs with overlapping GO terms are shown in Table 3 and Table 4. We also include AUPRC for these comparisons in Tables 5 and 6.

**Table 3.**
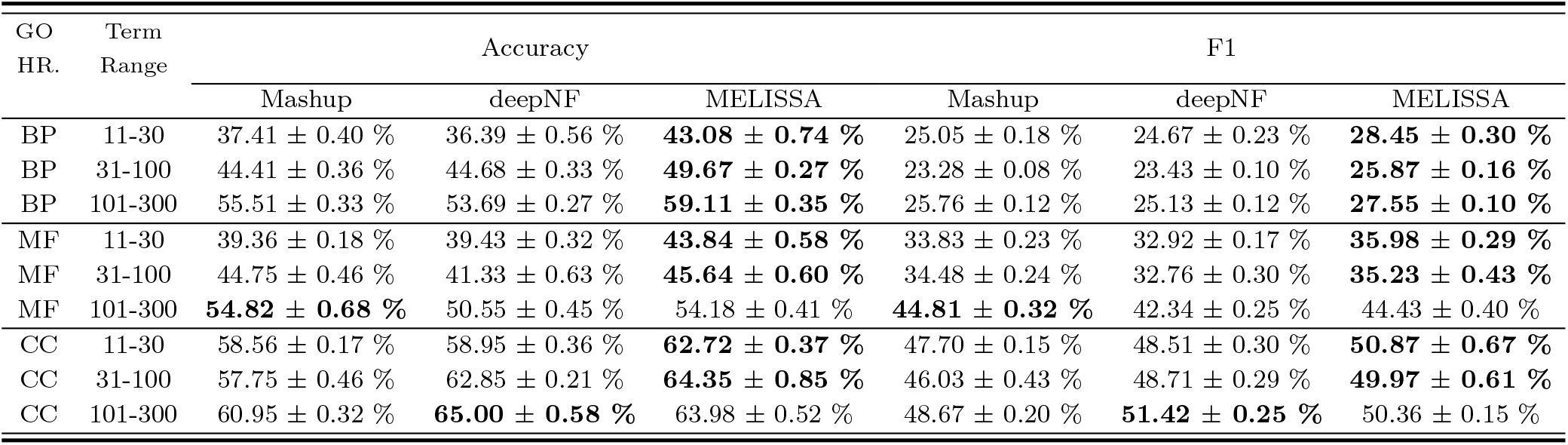
Function prediction performance on yeast PPI network with overlap included GO functional labels from the BP, MF, and CC hierarchies based on MELISSA, Mashup, and deepNF data features by majority voting with 10 nearest neighbors in 5-folds cross-validation. The embedding dimension is 400 for Mashup and MELISSA and the MELISSA parameters are: *w*^+^ = 1, *w*^−^ = −1, and 2 biclusters. The deepNF embedding dimension is 600. Results are averaged from 5 independent runs.

**Table 4.**
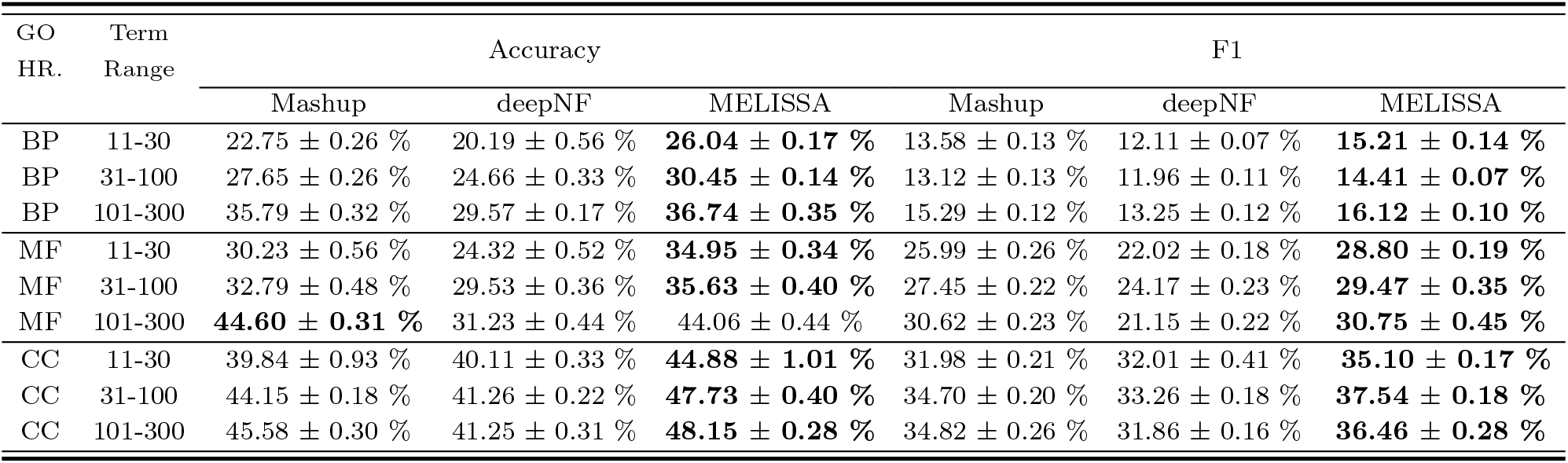
Function prediction performance on human PPI network with overlap included GO functional labels from the BP, MF, and CC hierarchies based on MELISSA, Mashup, and deepNF data features by majority voting with 10 nearest neighbors in 5-fold cross-validation. The embedding dimension is 400 and the MELISSA parameters are: *w*^+^ = 1, CL *w*^−^ = 1, and 2 biclusters. The deepNF embedding dimension is 1200. Standard deviation was calculated from 5 independent runs.

| GO<br>HR. | Term<br>Range | Accuracy |  |  | F1 |  |  |
| --- | --- | --- | --- | --- | --- | --- | --- |
|  |  | Mashup | deepNF | MELISSA | Mashup | deepNF | MELISSA |
| BP | 11-30 | 22.75 ± 0.26 % | 20.19 ± 0.56 % | **26.04 ± 0.17 %** | 13.58 ± 0.13 % | 12.11 ± 0.07 % | **15.21 ± 0.14 %** |
| BP | 31-100 | 27.65 ± 0.26 % | 24.66 ± 0.33 % | **30.45 ± 0.14 %** | 13.12 ± 0.13 % | 11.96 ± 0.11 % | **14.41 ± 0.07 %** |
| BP | 101-300 | 35.79 ± 0.32 % | 29.57 ± 0.17 % | **36.74 ± 0.35 %** | 15.29 ± 0.12 % | 13.25 ± 0.12 % | **16.12 ± 0.10 %** |
| MF | 11-30 | 30.23 ± 0.56 % | 24.32 ± 0.52 % | **34.95 ± 0.34 %** | 25.99 ± 0.26 % | 22.02 ± 0.18 % | **28.80 ± 0.19 %** |
| MF | 31-100 | 32.79 ± 0.48 % | 29.53 ± 0.36 % | **35.63 ± 0.40 %** | 27.45 ± 0.22 % | 24.17 ± 0.23 % | **29.47 ± 0.35 %** |
| MF | 101-300 | **44.60 ± 0.31 %** | 31.23 ± 0.44 % | 44.06 ± 0.44 % | 30.62 ± 0.23 % | 21.15 ± 0.22 % | **30.75 ± 0.45 %** |
| CC | 11-30 | 39.84 ± 0.93 % | 40.11 ± 0.33 % | **44.88 ± 1.01 %** | 31.98 ± 0.21 % | 32.01 ± 0.41 % | **35.10 ± 0.17 %** |
| CC | 31-100 | 44.15 ± 0.18 % | 41.26 ± 0.22 % | **47.73 ± 0.40 %** | 34.70 ± 0.20 % | 33.26 ± 0.18 % | **37.54 ± 0.18 %** |
| CC | 101-300 | 45.58 ± 0.30 % | 41.25 ± 0.31 % | **48.15 ± 0.28 %** | 34.82 ± 0.26 % | 31.86 ± 0.16 % | **36.46 ± 0.28 %** |

**Table 5.** AUPRC for function prediction on yeast PPI network with overlap included and overlap removed GO functional labels from the BP, MF, and CC hierarchies based on MELISSA, Mashup, and deepNF data features by majority voting with 10 nearest neighbors in 5-folds cross-validation. The embedding dimension is 400 for Mashup and MELISSA and the MELISSA parameters are: *w*^+^ = 1, *w*^−^ = −1, and 2 biclusters. The deepNF embedding dimension is 600. Results are averaged from 5 independent runs.

| GO<br>HR. | Term<br>Range | AUPRC (overlap removed) |  |  | AUPRC (overlap included) |  |  |
| --- | --- | --- | --- | --- | --- | --- | --- |
|  |  | Mashup | deepNF | MELISSA | Mashup | deepNF | MELISSA |
| BP | 11-30 | 37.91 ± 0.45 % | 36.98 ± 0.64 % | **43.32 ± 0.65 %** | 28.30 ± 0.25 % | 28.71 ± 0.43 % | **31.35 ± 0.48 %** |
| BP | 31-100 | 46.85 ± 0.35 % | 46.45 ± 0.65 % | **50.76 ± 0.48 %** | 34.47 ± 0.25 % | 34.05 ± 0.10 % | **36.11 ± 0.21 %** |
| BP | 101-300 | 59.99 ± 0.46 % | 58.85 ± 0.37 % | **63.45 ± 0.56 %** | 44.55 ± 0.34 % | 43.73 ± 0.15 % | **45.72 ± 0.27 %** |
| MF | 11-30 | 41.47 ± 0.83 % | 42.10 ± 0.54 % | **47.77 ± 0.52 %** | 31.41 ± 0.47 % | 30.46 ± 0.23 % | **34.05 ± 0.30 %** |
| MF | 31-100 | 45.00 ± 0.26 % | 42.57 ± 0.55 % | **46.21 ± 1.01 %** | 34.46 ± 0.24 % | 32.41 ± 0.18 % | **34.85 ± 0.19 %** |
| MF | 101-300 | 56.26 ± 0.64 % | 51.01 ± 0.41 % | **56.90 ± 0.87 %** | 41.98 ± 0.30 % | 40.09 ± 0.37 % | **42.19 ± 0.48 %** |
| CC | 11-30 | 62.74 ± 0.37 % | 63.95 ± 0.42 % | **64.51 ± 1.04 %** | 39.31 ± 0.12 % | **40.74 ± 0.17 %** | 40.08 ± 0.54 % |
| CC | 31-100 | 60.86 ± 0.80 % | 65.54 ± 0.13 % | **66.55 ± 0.55 %** | 41.27 ± 0.14 % | 42.79 ± 0.18 % | **43.87 ± 0.07 %** |
| CC | 101-300 | 71.49 ± 0.27 % | **76.00 ± 0.39 %** | 71.59 ± 0.93 % | 47.07 ± 0.14 % | **47.93 ± 0.20 %** | 45.89 ± 0.80 % |

**Table 6.** AUPRC for function prediction on human PPI network with overlap included and overlap removed GO functional labels from the BP, MF, and CC hierarchies based on MELISSA, Mashup, and deepNF data features by majority voting with 10 nearest neighbors in 5-folds cross-validation. The embedding dimension is 400 for Mashup and MELISSA and the MELISSA parameters are: *w*^+^ = 1, *w*^−^ = −1, and 2 biclusters. The deepNF embedding dimension is 1200. Results are averaged from 5 independent runs.

| GO<br>HR. | Term<br>Range | AUPRC (overlap removed) |  |  | AUPRC (overlap included) |  |  |
| --- | --- | --- | --- | --- | --- | --- | --- |
|  |  | Mashup | deepNF | MELISSA | Mashup | deepNF | MELISSA |
| BP | 11-30 | 12.74 ± 0.17 % | 10.19 ± 0.10 % | **14.61 ± 0.11 %** | 7.64 ± 0.11 % | 5.90 ± 0.09 % | **9.42 ± 0.09 %** |
| BP | 31-100 | 16.22 ± 0.23 % | 13.37 ± 0.10 % | **19.14 ± 0.14 %** | 10.48 ± 0.12 % | 8.97 ± 0.08 % | **12.62 ± 0.08 %** |
| BP | 101-300 | 30.08 ± 0.09 % | 25.61 ± 0.13 % | **33.70 ± 0.11 %** | 15.69 ± 0.16 % | 12.82 ± 0.12 % | **17.64 ± 0.11 %** |
| MF | 11-30 | 22.97 ± 0.35 % | 20.49 ± 0.33 % | **26.62 ± 0.23 %** | 20.96 ± 0.26 % | 15.29 ± 0.19 % | **23.85 ± 0.14 %** |
| MF | 31-100 | 23.87 ± 0.28 % | 19.61 ± 0.36 % | **26.69 ± 0.27 %** | 22.60 ± 0.24 % | 18.75 ± 0.15 % | **24.61 ± 0.22 %** |
| MF | 101-300 | 42.10 ± 0.12 % | 28.18 ± 0.14 % | **43.30 ± 0.26 %** | **30.81 ± 0.19 %** | 21.10 ± 0.31 % | 30.44 ± 0.49 % |
| CC | 11-30 | 38.49 ± 0.55 % | 39.28 ± 0.38 % | **44.09 ± 0.40 %** | 32.91 ± 0.42 % | 34.25 ± 0.39 % | **37.66 ± 0.40 %** |
| CC | 31-100 | 41.25 ± 0.22 % | 40.47 ± 0.22 % | **45.61 ± 0.14 %** | 35.04 ± 0.19 % | 35.25 ± 0.10 % | **39.84 ± 0.19 %** |
| CC | 101-300 | 44.13 ± 0.21 % | 40.70 ± 0.16 % | **47.07 ± 0.11 %** | 34.75 ± 0.16 % | 32.54 ± 0.23 % | **38.22 ± 0.22 %** |

As shown in Tables 5 and 6, as well as the pairwise differences between Tables 1 and 3 and Tables 2 and 4, the overall performance drops after re-introducing the overlapping GO terms in both yeast and human experiments. Even though MELISSA’s parameters were chosen for the non-overlapping GO term setting, we find that even with the overlapping GO terms retained, MELISSA outperforms Mashup and deepNF in most trials.

## Discussion

The results in Section 3 demonstrate that incorporating annotation information at an earlier stage can lead to more informative network embeddings. Augmenting the networks with cluster nodes and introducing ML and CL constraints in the embedding procedure introduces significant structures that are otherwise missed. This creates a more valuable and information-rich node embedding which yields performance gains in the functional enrichment of the local neighborhoods, as evidenced by the improvements in *k*NN-based functional classification. However, we note that Mashup improves performance by using more sophisticated downstream learning: namely, learning SVMs to discriminate functional classes. The SVM is expressive enough to overcome the generic embedding and performs better on our function prediction task than the Mashup or the Melissa embedding paired with *kNN*. However, the story is far from complete, and there is much room for exploring new techniques, especially given the wealth of work in semi-supervised methods.

One future direction is to push the annotations into the diffusion process for complete end-to-end utilization of all the information provided from the start. Many methods could be explored by modifying the topology of the networks using constraints. For example, changing the diffusion process to avoid transitioning directly between pairs of proteins with different functional labels. Other methods that could sparsify the networks or introduce new edges could also be investigated.

## Acknowledgements

Thanks to the T-Tripods graduate fellows and the T-Tripods summer reading group on graphs and networks members at Tufts. Thanks especially to T-Tripods graduate fellow Matthew Werenski who made important suggestions in the early stages of this project. This research was supported by NSF grants DMS-1812503 (to L.C. and X.H.) and the Tufts T-Tripods Institute (NSF grant CCF-1934553) and NSF CC* grant 2018149.

## References

1. E. Bair. Semi-supervised clustering methods. Wiley Interdisciplinary Reviews: Computational Statistics, 5(5):349–361, 2013.

2. M. Belkin, P. Niyogi, and V. Sindhwani. Manifold regularization: A geometric framework for learning from labeled and unlabeled examples. Journal of machine learning research, 7(11), 2006.

3. A. Ben-Hur, C. S. Ong, S. Sonnenburg, B. Schölkopf, and G. Rätsch. Support vector machines and kernels for computational biology. PLoS computational biology, 4(10):e1000173, 2008.

4. G. Berriz, O. King, B. Bryant, C. Sander, and F. Roth. Characterizing gene sets with FuncAssociate. Bioinformatics, 19(18):2502–2504, 2003.

5. M. Cao, C. M. Pietras, et al. New directions for diffusion-based prediction of protein function: incorporating pathways with confidence. Bioinformatics, 30:i219–i227, 2014.

6. M. Cao, H. Zhang, J. Park, N. M. Daniels, M. E. Crovella, L. J. Cowen, and B. Hescott. Going the distance for protein function prediction: a new distance metric for protein interaction networks. PloS one, 8(10):e76339, 2013.

7. H. Cho, B. Berger, and J. Peng. Compact integration of multinetwork topology for functional analysis of genes. Cell systems, 3(6):540–548, 2016.

8. G. O. Consortium. The gene ontology resource: 20 years and still GOing strong. Nucleic acids research, 47(D1):D330–D338, 2019.

9. I. S. Dhillon, S. Mallela, and D. S. Modha. Informationtheoretic coclustering. In Proceedings of the Ninth ACM SIGKDD International Conference on Knowledge Discovery and Data Mining, KDD ‘03, page 89–98, New York, NY, USA, 2003. Association for Computing Machinery.

10. A. M. Edwards, B. Kus, R. Jansen, D. Greenbaum, J. Greenblatt, and M. Gerstein. Bridging structural biology and genomics: assessing protein interaction data with known complexes. TRENDS in Genetics, 18(10):529–536, 2002.

11. A. Franceschini, D. Szklarczyk, S. Frankild, M. Kuhn, M. Simonovic, A. Roth, J. Lin, P. Minguez, P. Bork, C. vonMering, and L. Jensen. STRING v9.1: protein-protein interaction networks, with increased coverage and integration. Nucleic Acids Res., 41:D808–815, 2013.

12. G. Fu, J. Wang, B. Yang, and G. Yu. NegGOA: negative GO annotations selection using ontology structure. Bioinformatics, 32(19):2996–3004, 2016.

13. J. Gallier. Spectral theory of unsigned and signed graphs. applications to graph clustering: a survey. arXiv preprint arXiv:1601.04692, 2016.

14. V. Gligorijević, M. Barot, and R. Bonneau. deepnf: deep network fusion for protein function prediction. Bioinformatics, 34(22):3873–3881, 2018.

15. A. Grover and J. Leskovec. node2vec: Scalable feature learning for networks. In Proc. 22nd ACM SIGKDD, pages 855–864. ACM, 2016.

16. Y. P. Hou. Bounds for the least Laplacian eigenvalue of a signed graph. Acta Mathematica Sinica, 21(4):955–960, 2005.

17. S. Kohler, S. Bauer, D. Horn, and P. N. Robinson. Walking the interactome for prioritization of candidate disease genes. Am J Hum Genet., 82:949–958, 2008.

18. J. Kunegis, S. Schmidt, A. Lommatzsch, J. Lerner, E. W. De Luca, and S. Albayrak. Spectral analysis of signed graphs for clustering, prediction and visualization. In Proceedings of the 2010 SIAM International Conference on Data Mining, pages 559–570. SIAM, 2010.

19. C.-S. Liao, K. Lu, M. Baym, R. Singh, and B. Berger. IsoRankN: spectral methods for global alignment of multiple protein networks. Bioinformatics, 25(12):i253–i258, 2009.

20. V. A. Padilha and R. J. Campello. A systematic comparative evaluation of biclustering techniques. BMC bioinformatics, 18(1):1–25, 2017.

21. A. Ruepp, A. Zollner, D. Maier, K. Albermann, J. Hani, M. Mokrejs, I. Tetko, U. Guldener, G. Mannhaupt, M. Munsterkotter, and H. W. Mewes. The FunCat, a functional annotation scheme for systematic classification of proteins from whole genomes. Nucleic Acids Res, 32:5529–5545, 2004.

22. the Gene Ontology Consortium. Gene Ontology: tool for the unification of biology. Nature Genetics, 25(1):25–29, 2000. http://www.geneontology.org.

23. H. Tong, C. Faloutsos, and J. Pan. Fast random walk with restart and its applications. In Sixth International Conference on Data Mining (ICDM’06), pages 613–622, 2006.

24. P. Vincent, H. Larochelle, I. Lajoie, Y. Bengio, P.-A. Manzagol, and L. Bottou. Stacked denoising autoencoders: Learning useful representations in a deep network with a local denoising criterion. Journal of machine learning research, 11(12), 2010.

25. A. Warwick Vesztrocy and C. Dessimoz. Benchmarking gene ontology function predictions using negative annotations. Bioinformatics, 36(Supplement 1):i210–i218, 2020.

26. J. Weston, F. Ratle, and R. Collobert. Deep learning via semi-supervised embedding. In Proceedings of the 25th international conference on Machine learning, pages 1168–1175, 2008.

27. D. Zhang, Z.-H. Zhou, and S. Chen. Semi-supervised dimensionality reduction. In Proceedings of the 2007 SIAM International Conference on Data Mining, pages 629–634. SIAM, 2007.

28. X. Zhu, Z. Ghahramani, and J. D. Lafferty. Semi-supervised learning using gaussian fields and harmonic functions. In Proceedings of the 20th International conference on Machine learning (ICML-03), pages 912–919, 2003.

